# The Critical Role of *Pdyn*-Lineage Enteric Neurons in Colonic Motility and Visceral Interoception

**DOI:** 10.64898/2026.07.21.739264

**Authors:** Daniel Joseph Verbaro, Jun-Nan Li, Prashant Gupta, Bradley Jacobo, Juliet Mwirigi, Adam Dourson, Robert W Gereau

## Abstract

Summary/Abstract

Sensory neurons play well defined roles in the regulation of intestinal motility, digestion, and interoception, but transcriptional dissection of intrinsic and extrinsic sensory neurons innervating the intestines has been challenging because these cells share many genetic markers. However, we cross-referenced transcriptional profiles of intrinsic and extrinsic intestinal neurons and the cells that surround them and identified *Pdyn,* the gene encoding Prodynorphin, as a marker of putative sensory enteric neurons of the mouse intestines. In a *Pdyn* lineage-reporter mouse, we identified labeled cells in the myenteric and submucosal plexuses of the large intestine, in contrast to their sparse presence in the dorsal root or nodose ganglia. In dissociated cell culture, these neurons mostly display a phasic firing pattern, discharging one or two action potentials at the onset of a depolarizing current pulse, followed by a prompt cessation of firing despite continued current injection, as assessed by whole-cell patch-clamp recordings. Optogenetic activation of *Pdyn*-lineage neurons propels stool in *ex vivo* colons, and in untethered and mobile mice, optogenetic stimulation of the proximal colon induces freezing behaviors and orbital tightening, suggesting interoceptive behaviors, without significant stool output differences. Together, these findings suggest that activation of *Pdyn*-lineage enteric neurons regulates motility and visceral interoception.

## Introduction

The gastrointestinal tract detects sensory stimuli important for motility and digestion of nutrients, and interoception^1, 2^. These stimuli are detected by an intricate network of neurons whose soma reside within or outside of the intestinal wall^3^. Neurons within the wall comprise the enteric nervous system (ENS), whereas those with soma outside the intestinal wall include the vagal and dorsal root ganglia (DRG) neurons^3^. Studying the individual contributions of these neurons to gastrointestinal function has been difficult because of their close anatomical relationships and extensively overlapping gene expression profiles. Murine intersectional genetic targeting techniques, such as combinatorial genetic recombination via multiple recombinases^4, 5^ and locally restricted genetic recombination via virally expressed recombinases^1, 6^, have provided valuable information about the location and function of transcriptionally defined enteric and DRG neurons in mice. However, these techniques have not completely characterized the function and electrophysiological properties of all the transcriptionally defined neuron populations that express human analogous genes implicated in intestinal function.

Recent data demonstrate substantial heterogeneity in transcriptional profiles of enteric neurons, including populations considered to be sensory neurons based on their expression of the genes encoding calcitonin gene-related peptide and calbindin^7, 8^. Some of these transcriptionally defined sensory enteric neurons have been functionally investigated^9^, but the role of transcriptionally defined enteric neurons that express the mechanoreceptor *Piezo2* remains elusive^10^. This subpopulation is of particular interest because broadly targeting *Piezo2* in the colon induces pain-like behaviors in mice^1^, but it is not known if enteric neurons expressing *Piezo2* are sufficient for this response. *Piezo2* is expressed in myocytes, fibroblasts, enteroendocrine cells, glia and lymphatic and vascular endothelial cells^7^, and prior studies of *Piezo2*-expressing cells in the colon used a ubiquitous promoter^1^. Nonetheless, there are sufficient data to identify sensory neuron clusters that express *Piezo2*, which are noteworthy because in addition to *Piezo2*^10–12^, these cells (termed “putative sensory population 4 (PSN4)” and “ENS12”) express *Bdnf*^13–15^ and *Chat*^16, 17^, which are relevant to human somatic and abdominal pain. Although not expressed in ENS12, *Hes6*^18^, *Trpm3*^19–21^, *Cnr1*^22, 23^, and *Oprm1*^24^ are also implicated in pain and expressed in the PSN4 subset. Whether these transcriptionally defined populations are identical remains unclear, as they did not align after harmonization of the sequencing datasets^6^. Moreover, comprehensive mapping of the ENS did not completely describe these neurons^6^, which may highlight the limitation of finding rarer populations with fluorescence-activated cell sorting (FACS).

To study the putative sensory cluster PSN4, we hypothesized that these neurons could be targeted with a genetic reporter relying on a single genetically driven recombinase, without the need of multiple recombinases or viral-driven recombinases. To identify such a target, we cross-referenced transcriptional profiles of enteric neurons^7^ and dorsal root ganglia (DRG) neurons^25^. Here, we report the differential expression of *Pdyn,* the gene encoding Prodynorphin, and identified neurons that express this gene within their lineage as mediators of intestinal motility and pain perception in an informative murine model.

## Results

There is overlap in transcriptional signatures between enteric neurons, neurons of the dorsal root ganglia (DRG), and surrounding non-neuronal intestinal cells complicating cellular functional studies utilizing genetic targeting^6–8, 25^. However, the available RNA-sequencing datasets of the intestines and DRG provide an opportunity to highlight putative selective genetic markers for studying subsets of enteric neurons. To investigate enteric neurons that express genes relevant to human gastrointestinal motility disorders and pain such as *Piezo2*, *Bdnf*, *Chat*, *Hes6*, and *Oprm1*, we cross-referenced transcriptional profiles from mouse enteric neurons with neurons of the dorsal root ganglia and surrounding non-neuronal cells^7, 25^. One putative gene for targeting these enteric neurons was *Pdyn*, encoding Prodynorphin (Figure 1A) given that its expression was specific to enteric neurons in these RNA-sequencing datasets. With this specificity, we hypothesized that *Pdyn* promoter/enhancer element expression could be used to target a subset of enteric neurons expressing genes relevant to human gastrointestinal disorders.

**Figure 1.**
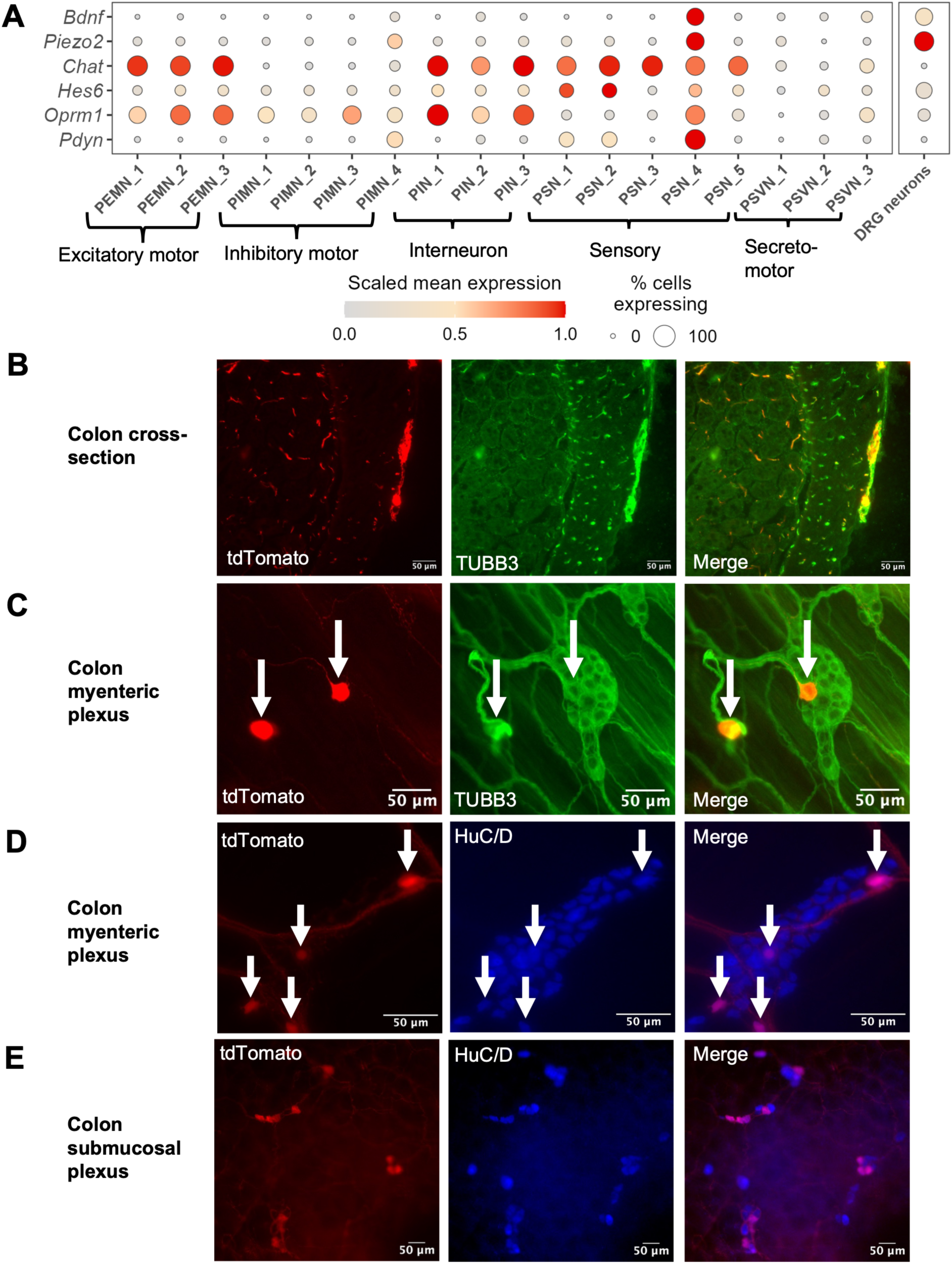
*Pdyn* expression marks a subset of myenteric neurons. **A**. Heat map showing expression of *Bdnf, Piezo2, Chat, Hes6, Oprm1*, and *Pdyn* transcripts from mouse large bowel neurons obtained from Broad Institute Single Cell portal^7^. **B**. Heat map showing expression of *Bdnf, Piezo2, Chat, Hes6, Oprm1*, and *Pdyn* transcripts from dorsal root ganglia of mouse from NIH Precision Human Pain Network’s Pain-seq Harmonized DRG sequencing data^25^. **C**. Representative images of cryosections of colon from offspring of *Pdyn*-cre mice bred with Ai14 mice (*Pdyn*^tdTomato^) showing neuronal tubulin protein staining with anti-TUBB3 in green. **D**. Representative images of whole mount longitudinal muscle-myenteric plexus (LMMP) preps from *Pdyn*^tdTomato^ mice (tdtomato: Red) stained with anti-TUBB3 (green). **E**. Representative images of whole mount longitudinal muscle-myenteric plexus (LMMP) preps from *Pdyn*^tdTomato^ mice (tdtomato: Red) showing staining of neuronal nuclei with anti-HuC/D (blue). **F**. Representative images of whole mount submucosal plexus preps from *Pdyn*^tdTomato^ mice stained with anti-HuC/D (blue).

Next, we generated a *Pdyn* reporter mouse by crossing the *Pdyn*-IRES-Cre mouse line to the fluorescent reporter mouse line (Ai14)^26, 27^. Therefore, tdTomato signal marks cells expressing *Pdyn* within their lineage, which mice are designated as *Pdyn*^tdTomato^. In cross-sections of the mouse colon, tdTomato was observed in the myenteric layer and interspersed throughout the submucosa (Figure 1B), and these projections co-localized with the neuronal marker, β-III Tubulin. There was a lack of tdTomato signal in non-neuronal cells. In longitudinal muscle/myenteric plexus and submucosal whole mounts, tdTomato co-localized with markers for neurons such as β-III Tubulin and HuC/D (Figure 1B-E). These data suggest that lineage expression of *Pdyn* selectively labels neurons within the mouse colon without the need for intra-wall injection or multiple recombinases for labeling. Further supporting the specificity of *Pdyn* for enteric neurons of the intestines, we found sparse labeling of neurons within the dorsal root ganglia at vertebral levels (T13-S2) known to send projections to the mouse large intestine (Figure S1A). There was rare expression within the nodose ganglia, which contain soma of afferent neurons of the vagus nerve (Figure S1B). The total number of labeled neurons in the DRG and nodose are shown in Figure S1C. The total number of DRG and nodose cells that express Pdyn within their lineage is rare because the total number of neurons per ganglion is on the order of 10^3^. The *Pdyn* promoter/enhancer element expression, therefore, reliably and selectively labels a subset of enteric neurons found within the large intestine allowing for identification for electrophysiological and *in situ* functional measurements via optogenetics with minimal off-target effects.

We next investigated the electrophysiological properties of *Pdyn*-expressing cells from the mouse colon. Myenteric neurons were cultured from *Pdyn* lineage-reporter mice as previously described^28^. Cells labeled with tdTomato remained identifiable to at least day 5 of culture and expressed neuronal protein, β-III Tubulin (Figures 2A). As noted by β-III Tubulin, labeled cells displayed multiple axonal projections (arrows in Figure 2A), which is a characteristic of dogiel type II cells^8, 29^. This classification had been proposed to identify intrinsic primary afferent neurons (IPANs)^30^. There were some cultured cells that did not form any projections. The lineage reporter allowed identification and targeting for whole-cell patch-clamp recordings as shown in Figure 2B. Of the recorded cells that express *Pdyn* in their lineage, the average major axis diameter was 14.4µm ± 0.7 µm and average minor axis diameter was 11.2 ± 0.6 µm (Figure 2C). Prior to injecting currents, labeled cells were found to have an average resting membrane potential of -45.5 ± 1.0 mV with an average input resistance of 999.5 ± 112.4 MΩ (Figures 2D-E). With increasing current injection up to the maximum of 225 pA used in this study, we found that most labeled cells exhibited a phasic firing pattern, generating 1-2 action potentials at the onset of a depolarizing current pulse (Figures 2F-G). Of all recorded cells, there was 1 cell that fired more than 2 action potentials with increased current (Figures 2F-G). The I_threshold_, or lowest current to elicit an action potential, was 27.0 ± 5.0pA (Table S1). About 90% of the cells elicited a slow after-hyperpolarization (Figure 2H), which is characteristic of AH (after-hyperpolarization)-type enteric neurons^9, 31, 32^. About 10% of the cells elicited an inflection point in the first derivative of the action potential (Figure 2I), which most had only a fast phase to repolarization. The slower phase of repolarization has been noted in AH type neurons^33^, but this feature was rarely noted in neurons displaying AH-type properties. The action potential half-width was 3.9 ± 1.5 ms and amplitude was 45.7 ± 3.7 mV (Table S1). Other intrinsic properties of *Pdyn*-expressing cells are summarized in Tabel 1. From these data, we found that cultured colonic cells that express *Pdyn* in their lineage generate action potentials with current injection. Myenteric neurons that express Pdyn in their lineage display some electrophysiological properties akin to sensory or intrinsic primary afferent neurons.

**Figure 2.**
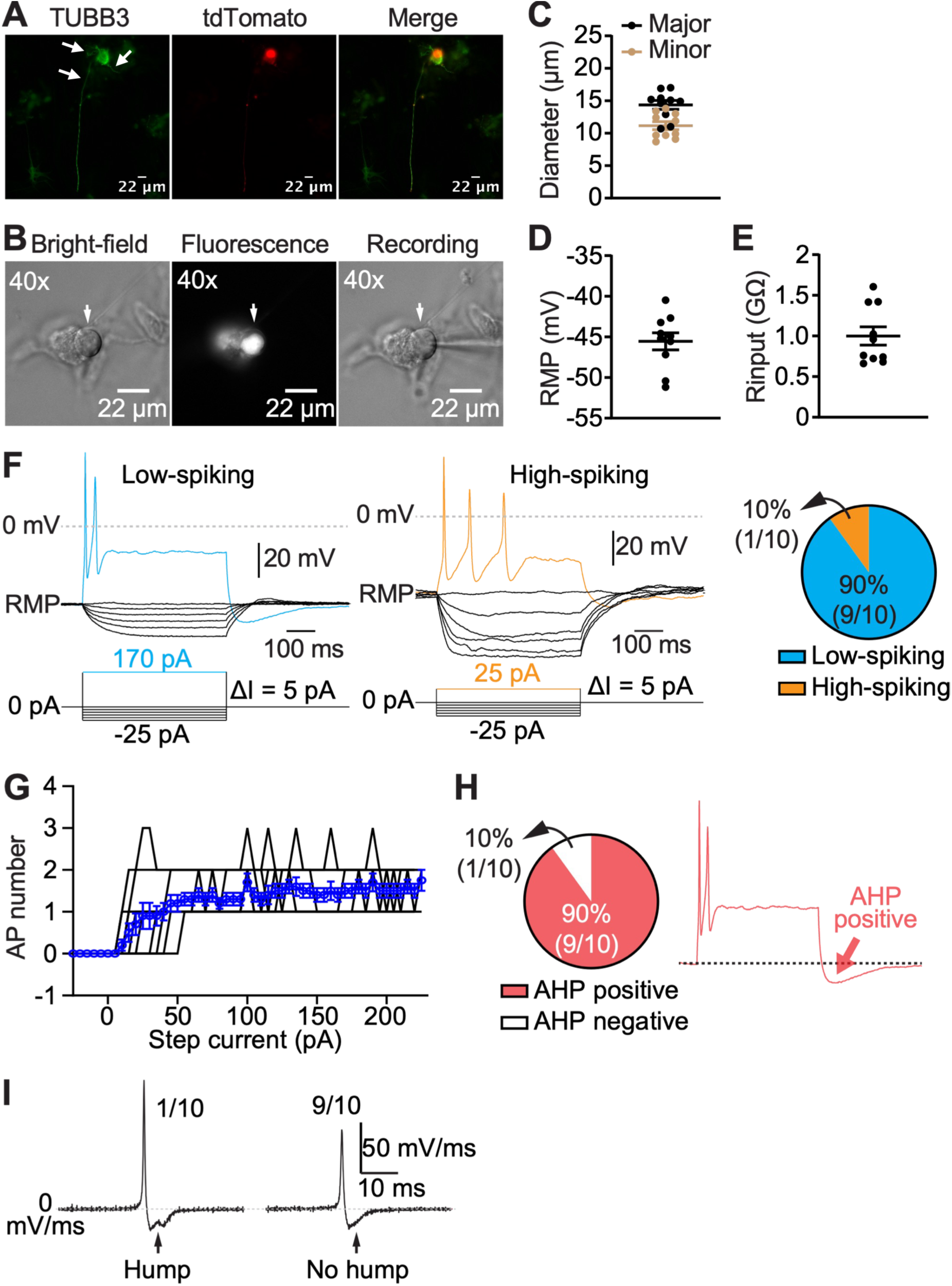
Electrophysiological properties of cultured *Pdyn*-expressing myenteric neurons. **A**. Representative images of cultured labeled myenteric cells from *Pdyn*^tdTomato^ mice stained with anti-TUBB3 antibody (green). **B**. Representative images of cultured myenteric cells from *Pdyn*^tdTomato^ mice during whole-cell patch-clamp recordings showing identification and targeting of labeled cells. **C**. Soma dimensions of *Pdyn*-expressing myenteric cells, showing major (black) and minor (brown) axes. **D**. Resting membrane potential and **E**. the input resistance of *Pdyn*-expressing myenteric neurons. **F**. Representative membrane potential traces (top) in response to stepped current injections (bottom). The pie chart (right) indicates the relative distribution of low-spiking (90% blue) and high-spiking (10% orange) populations within the sample (n=10). **G**. Input-output curve showing action potential counts per current step for 10 individual cells (black lines) and the group average (blue line; error bars indicate SEM). **H**. Representative trace of a neuron (right) showing the deflection below the resting membrane potential, which is the after-hyperpolarization (AHP), and the percentage of cells exhibiting an AHP are shown in the pie chart (left). **I**. Representative first derivative of electrophysiological traces showing an example cell that exhibited an inflection in the first derivative and a cell that did not.

To study the functional outcome of activating these enteric neurons, we utilized specific activation of these cells via optogenetics through genetically driven channelrhodopsin-2 (ChR2) by crossing the *Pdyn*-IRES-cre mouse to the ROSA26^LSL-ChR2-EYFP^ mouse^34^. This strategy would also avoid off-target effects of *Pdyn*-expressing neurons in the brain and spinal cord. Offspring express the light activated ion channel, ChR2-EYFP fusion protein, in cells that express *Pdyn* in their lineage, which mice are designated as *Pdyn*^ChR2-EYFP^. To determine the optimal parameters for subsequent *ex vivo* colons and *in vivo* optogenetic stimulation, we validated the expression and functional frequency limits of ChR2 in cultured *Pdyn*-expressing cells. First, we determined whether 470nm light activated *Pdyn*-expressing cells by performing optogenetic stimulation on cultured enteric *Pdyn*-expressing cells. EYFP expressing cells were identifiable after several days of culture from dissociated myenteric plexus preparations (Figure S2A-C). Upon pulses of 470nm light, actions potentials were observed. We observed reliable action potential generation (≥80% success rate) in response to light pulses at frequences up to 10hz (20 ms pulse width, 20 mW/mm^2^) during 30 s stimulation (Figures S2D-E). At higher frequencies, action potential generation became less reliable on a per-pulse basis (Figures S2D-F). A comparable success rate of light-evoked action potentials was observed after a 2 min recovery periods during a second round of stimulation (red line of Figure S2D). These data confirm that in these mice light activates neurons that expressed *Pdyn* within their lineage.

Next, we performed optogenetic studies on *ex vivo* colons from *Pdyn*^ChR2-EYFP^ mice to understand the impact of activity in these neurons on natural stool pellet movement. Upon activation of the proximal colon with 470nm light at 10Hz and light intensity 20mW/mm^2^ (these parameters were selected based on the reliable action potential generation *in vitro*), we noted anterograde movement of natural stool pellets. Movement initiated after the light stimulus occurred at a significantly higher frequency in colons expressing ChR2 in *Pdyn*-lineage cells compared to control colons that exhibit spontaneous pellet movement (Figure 3A-B). Natural stool pellets moved from the proximal to distal after light pulses in *Pdyn*^ChR2-EYFP^ colons supporting anterograde movement (Figure 3A). More propagating anterograde contractions were noted in colons expressing ChR2 compared to controls with optogenetic stimulation (Figure 3C). The time to contraction from onset of light stimulus was shorter in colons expressing ChR2 compared to spontaneous contractions observed in control colons, further supporting that light activation induces natural pellet movement (Figure 3D). These data suggest that enteric neurons that express *Pdyn* in their lineage play a role in the coordination of anterograde stool movement.

**Figure 3.**
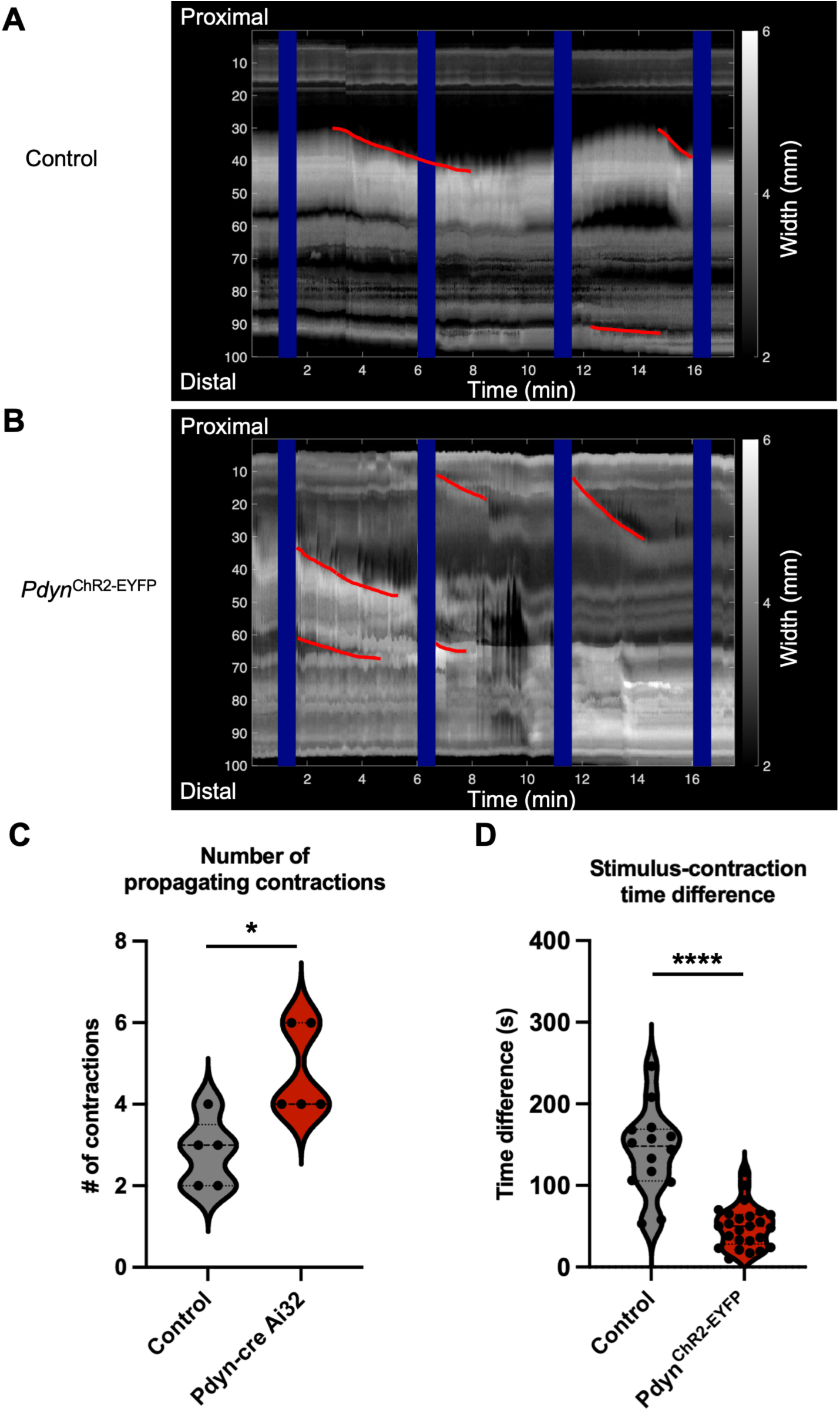
*Ex vivo* optogenetic stimulation of *Pdyn* expressing myenteric neurons caused increased contractility. Representative spatial-temporal maps generated from video recordings of *ex vivo* colons from control mice that harbor the LSL-ChR2-EYFP allele but lack the Cre recombinase (n=5 mouse colons) and (**A**) and Pdyn^ChR2-EYFP^ (n=5 mouse colons) (**B**) stimulated with 470nm light pulses with a frequency of 5Hz for 20 sec every 3 minutes and pulse width of 20ms. Blue lines indicate time of stimulation, and red lines highlight anterograde pellet movement. **C**. Number of stool propagating contractions of natural fecal pellets from control and *Pdyn*^ChR2-EYFP^ colons. Bars represent mean and SEM. p=0.0129 calculated by Welch’s two-tailed t test. **D**. Time difference between onset of stimulus and onset of stool propagating contraction from control and *Pdyn*^ChR2-EYFP^ colons. Bars represent mean and SEM. p value was <0.0001 when calculated by Welch’s two-tailed t test. **E**. Total number of stools during the stimulation was enumerated. Bars represent mean and SEM.

To determine the function of these enteric neurons in the mouse, we utilized *in vivo* optogenetics with biocompatible, wireless implantable LED’s^35^. The device body was implanted subcutaneously, and the LED was tunneled under the fascia superficial to the proximal colon. Mice were allowed to recover for at least 2 weeks prior to optogenetic stimulation. Mice were placed in a chamber lined by wires that control the LED parameters. Upon optogenetic stimulation, we observed that mice with ChR2 expression in enteric neurons that expressed *Pdyn* exhibited orbital tightening (Figure 4B-C) and freezing behaviors (Figure 4D) while control mice with LED implants did not. We did not observe significant differences in stool output between groups during stimulation (Figure 4E).

**Figure 4.**
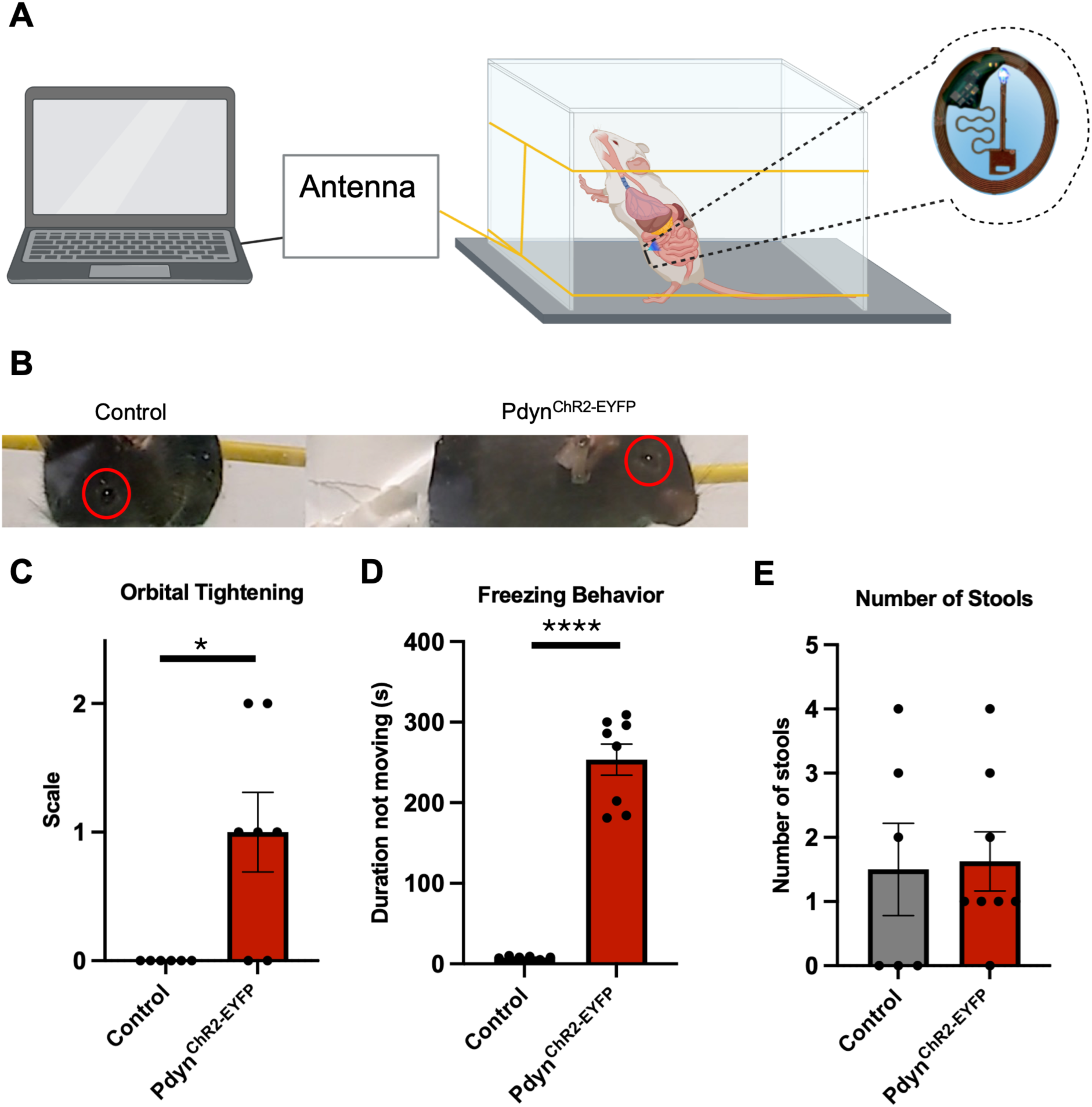
Optogenetic stimulation of proximal colonic *Pdyn* expressing neurons *in vivo*. **A**. Schematic of wireless optogenetic stimulation of the proximal colon of *Pdyn*-expressing intestinal cells. **B**. Representative still image of mouse eyes from video recordings of mice after LED pulses in control mice, which harbor the LSL-ChR2-EYFP allele but lack the Cre recombinase allele, and *Pdyn*^ChR2-EYFP^ mice. (**C**) Orbital tightening score as defined in mouse grimace scoring (p=0.0177), (**D**) Duration spent not moving (p<0.0001), and (**E**) number of stools produced was quantified after 5^th^ stimulation for control and *Pdyn*^ChR2-EYFP^ mice (p=0.7473). Two-tailed unpaired t-tests were performed. Schematic was generated in Biorender.com.

## Discussion

Considerable effort has been made to describe the anatomy and function of transcriptionally defined enteric neurons^5, 6, 35^; however, a complete functional and electrophysiological understanding of these neuronal subpopulations is still lacking. Such information has important implications for the development of new therapeutics targeting human gastrointestinal function and for predicting gastrointestinal side effects of therapeutics developed for non-GI indications. Most newly developed animal models for studying enteric nervous system function require extensive breeding or invasive abdominal surgery to achieve specific genetic manipulation of intestinal cells or neurons^1, 5, 6^. The extent to which abdominal surgery and intracolonic injections of the intestinal wall affect enteric neurons is not known. Therefore, in this study, we sought a less invasive way to target transcriptionally defined enteric neurons. Although our use of an implantable LED still requires abdominal surgery^35^, the fascial incision is considerably smaller, the surgical time is significantly shorter than the open laparotomy for intestinal AAV infection, and there is no manipulation of the bowel itself.

To identify a candidate gene for selective targeting of enteric neurons, we utilized ribosome-bound RNA and mining rare cells sequencing (MIRACL-seq) data^7^ and dorsal root ganglia RNA sequencing data^25^. Another study that transcriptionally defined enteric neurons utilized FACS-based neuronal purification^6^, but the diversity of enteric neuron subtypes was smaller compared with MIRACL-seq. Although rare neuronal populations were found in mice, these cells express many genes relevant to human disease. In this context, the *Pdyn* mouse models could be used to study such the function of Piezo2 in the enteric nervous system, whose role has been elusive to date^2, 10^. Mechanosensation is likely critical within the enteric nervous system itself, not only in dorsal root ganglia that innervate the intestines or enteroendocrine cells. Previous studies have demonstrated that optogenetic activation of the broad population of *Piezo2*-expressing cells of the distal colon and rectum elicits a visceromotor response^1^, however, the specific cellular contributors to this response remain to be fully elucidated. *Piezo2* is also expressed in the nodose ganglia of the vagus nerve^36^, dorsal root ganglia^10^, and intestinal enteroendocrine cells^37^, all of which contribute to visceral hypersensitivity and pain. Viral targeting has not yet overcome this limitation^1^. The promotors commonly used to drive expression in AAV- based approach, such as *Syn1* and *Eef1a1,* are not unique to enteric neurons and are also expressed in enteroendocrine cells^7^, raising the possibility of off-target effects. In contrast, our mouse model targets enteric neurons while avoiding off-target activation in non-neuronal colonic cells. Intriguingly, *Pdyn*, the gene encoding Prodynorphin, was specific to neurons in the mouse intestine, with no expression detected in surrounding intestinal cells and only rare expression in the dorsal root ganglia. A *Pdyn* lineage reporter mouse labeled cellular populations consistent with those identified in RNA-sequencing datasets from adult mouse dorsal root ganglia and enteric neurons. This concordance suggests that the common progenitor of enteric and DRG neurons is unlikely to express *Pdyn*, given the rare expression observed in DRG neurons.

Dynorphin was observed in the guinea pig enteric nervous system and was assigned to myenteric inhibitory circular muscle motor neurons and submucosal cholinergic secretomotor/ vasodilator neurons and cholinergic secretomotor (non-vasodilator) neurons, based on co-localization of vasoactive intestinal peptide (VIP) and galanin (GAL)^38^. Most cultured myenteric neurons that expressed *Pdyn* in their lineage exhibited a prolonged after-hyperpolarization within their electrophysiological properties, a feature characteristic of intrinsic primary afferent neurons (IPANs) in both guinea pig and mouse, but not of motor neurons and interneurons^31, 39, 40^. Although dissociation and culture likely alter some electrophysiological features, most of these neurons display AH-type properties in culture, suggesting that they may correspond to an IPAN or intestinal intrinsic sensory neuron population. Supporting cross-species conservation, human enteric neuron sequencing identified *PDYN*-expressing cells that clustered predominantly with afferent neurons of the enteric nervous system^41^.

Similar electrophysiological properties have been observed in intestinal neurons expressing *Cdh6*. However, optogenetic activation of *Cdh6*-expressing cells, which includes intestinal smooth muscle, results in retrograde colonic motor complexes^9^. These findings highlight the heterogeneity of functional outcomes among neurons with overlapping electrophysiological profiles but distinct transcriptional profiles, underscoring the importance of genetic approaches to dissecting enteric neuronal function. Additionally, the connectivity between transcriptionally defined enteric neurons is not known, and the enteric nervous system connectome is likely to play a critical role in functional outcomes such as motility. New genetic mouse models will provide opportunities to elucidate connections between transcriptionally distinct enteric neurons.

Activation of *Pdyn*-expressing enteric neurons in freely moving mice resulted in profound activity change and orbital tightening, which is a feature of mouse grimace, a facial feature used to assess pain in mice^42^. Thus, these cells likely contribute to visceral interoception and may contribute to the sensation of pain. A recent study found that activating cholinergic enteric neurons resulted in increased visceromotor responses and spontaneous hunching, other surrogate readouts for visceral interoception or pain^1^. Since many transcriptionally distinct enteric neurons are cholinergic, whether all distinct cholinergic neuron clusters contribute to pain is unknown. Our data suggest that activation of a subset of enteric neurons are sufficient to cause pain-like behaviors.

Inflammation rendered cholinergic neuron activation aversive^1^, but the mechanism by which inflammation mediates this effect is unknown. How inflammation affects enteric neurons that express Pdyn in their lineage and whether they contribute to inflammatory intestinal pain is unknown. Intriguingly, enteric neurons that express *Pdyn* were also found to express cytokine receptor subunits such as IL1R1 and IFNγR1. The RNA datasets suggest that these cells might respond directly to proinflammatory cytokines, but functional outcomes of this signaling in transcriptionally defined enteric neurons is not known. Our work here sets up a model to begin to study these effects and mechanism behind which inflammation affects the intrinsic properties of transcriptionally defined enteric neurons that upon activation are sufficient for visceral interoception behaviors.

### Methods Animals

All experimental procedures were approved by the Institutional Animal Care and Use Committee (IACUC) of Washington University School of Medicine and conducted in accordance with the National Institutes of Health guidelines for the care and use of laboratory animals. Mice were house in standard 12-hour light/dark cycle and provided food and water ad libitum. Male and female mice between the ages of 8 and 14 weeks were used for all experiments. *Pdyn*-IRES-cre (B6.129S-*Pdyn*^tm1.1(cre)Mjkr^/LowlJ; strain #: 027958) mice, AI14-tdtomato reporter mice (B6.Cg-Gt(ROSA)26Sor^tm14(CAG–tdTomato)Hze^/J; Strain #: 007914), Ai32-channelrhodopsin-2/eYFP (B6.129S-Gt(ROSA)26Sor^tm32(CAG–COP4*H134R/EYFP)Hze^/J; strain #: 012569) and C57B6 mice were purchased from The Jackson Laboratory.

### Immunohistochemistry

Mice were anesthetized with isoflurane and then euthanized by decapitation. The abdomen was dissected, and the colon was transected distally to the cecum and proximally to the rectum. For whole mount preparation, the colon was flushed with ice cold phosphate buffered saline (PBS) and then cut along the mesenteric line. The tissue was splayed with the serosa side up. The longitudinal muscle/myenteric plexus was separated from the epithelial and submucosal layers of the intestine by gentle retraction. The tissue was pinned on a Sylgard dish and bathed in 4% paraformaldehyde for 30 minutes at room temperature. Tissues were dehydrated with 30% sucrose overnight.

All tissues were blocked in 1% BSA, 10% normal goat serum, and 0.5% triton X in PBS for 1 hour at room temperature while rocking. Tissues were washed with PBS three times. Primary antibodies were incubated overnight at 4°C while rocking. Tissues were washed three times with PBS. Secondary antibodies were incubated overnight at 4°C while rocking. Tissues were washed three times and then placed on microscope slide and then VECTASHIELD® Hardset mounting media with DAPI (Vector Laboratories, H-1400) was added to tissue before placing on coverslip.

For cryosection preparation, colons were dissected and flushed with PBS. Tissue was fixed for 40 minutes and then moved to 30% sucrose solution overnight. Then the tissues were placed in Optimal Cutting Temperature (O.C.T.) compound and flash frozen on dry ice. Tissues were sectioned on Leica cryostat and placed on microscope slides. Similar staining protocol was used for sections as done for whole mount. Fluorescent images were obtained on a Leica epifluorescence microscope.

Table of antibodies

| Name | Dilution | Manufacturer | Catalog # | Lot # |
| --- | --- | --- | --- | --- |
| Rabbit Anti-HuC/D-Alexafluor 647 | 1:500 | Abcam | AB237234 | 1083472-5 |
| Rabbit Anti-TUBB3 | 1:500 | Biolegend | 802001 | B410083 |
| Alexa Fluor 488 goat anti-Rabbit | 1:1000 | ThermoFisher Scientific | A-11008 | 23800031 |

For DRG isolation, the spinal column was removed and transected anteriorly and posteriorly. The T13 through S2 dorsal root ganglia were dissected with fine forceps. For nodose extraction, anterior incision of the neck was performed exposing the submandibular glands, which were retracted laterally. The sternocleidomastoid and omohyoid muscle were then retracted laterally and medially, respectively. The carotid sheath was identified and then vagus nerve was isolated. The nerve was traced rostrally until the nodose was observed and removed with fine forceps with gentle retraction. These ganglia were placed in 4% paraformaldehyde for 30 minutes at room temperature.

Tissues were dehydrated with 30% sucrose overnight. The next day, tissues were placed in OCT and immediately frozen. Tissues were cut with a 25µm thickness on a Leica cryostat and placed on a microscope slide. Slices were stained with VECTASHIELD® Hardset mounting media with DAPI.

### Myenteric Neuron Culture

Mice were anesthetized with isoflurane and then euthanized by decapitation. The abdomen was dissected, and the colon was transected distally to the cecum and proximally to the rectum. The colon was flushed with ice cold Hanks’ Balanced Salt Solution (HBSS) and then cut along the mesenteric line. The tissue was splayed with the serosal side up in HBSS. The longitudinal muscle/myenteric plexus was separated from the other layers of the intestine by gentle retraction. The tissue was cut into 1mm x 1mm pieces and transferred to tube containing HBSS. Tissue was centrifuged at 360g for 30 seconds and HBSS was removed. Next, warmed type 2 collagenase (Sigma C6885, Batch: 0000127207) at 1mg/ml in HBSS supplemented with Hepes and calcium was added. This was incubated for 15 minutes at 37°C. Tissue was gently inverted every 5 minutes. After incubation, the tissue was centrifuged for 2 minutes at 360g and supernatant was removed. Warmed 0.05% Trypsin (ThermoFisher Scientific) in HBSS was added to the tissue and incubated at 37°C for 10 minutes with gentle inversion every 5 minutes. Samples were centrifuged and trypsin solution was removed. Tissue was resuspended in neuronal media: Neurobasal A (Gibco), 1% penicillin/streptomycin (Corning, Corning, NY), B27, 5% FBS (Gibco, Grand Island, NY), Glutamax (Life Technologies, Carlsbad, CA), and 10ng/mL GDNF (Peprotech Lot: 0607421). Tissue was titurated with 1ml pipet about 20 times. Tissue was allowed to settle to bottom, and supernatant was passed through 40µm filter. Additional neuronal media was added to the tissue and again titurated through an 1mL pipet tip about 20 times. All solution was filtered through 40µm filter. Cells were centrifuged for 8 minutes at 360g. Cells were plated on PEI and laminin coated coverslips. Cells were flooded with media about 1-3 hours after plating. 50% of media was replaced every 3^rd^ day of culture.

### Whole-cell patch clamp recordings and optogenetics in dissociated cultured cells

Electrophysiological recordings of myenteric cells were performed between days 5 and 7 of culture via whole-cell patch clamp configuration. The coverslips were transferred to a recording chamber (Scientific Instruments) and perfused continuously (2.0mL/min) with bubbled (continuously aerated with 95% O_2_/5% CO_2_) Kreb’s solution (118mM NaCl, 4.6mM KCl, 1.3mM NaH_2_PO_4_, 1.2mM MgSO_4_, 25mM NaHCO_3_, 11mM glucose, and 2.5mM CaCl_2_, pH=7.3-7.4 and osmolarity: 305-310) driven by a mini-peristaltic pump II (Harvard Apparatus). tdTomato expressing- or EYFP expressing-cells were visualized with differential interference contrast optics on an upright microscope (BX50WI, Olympus). The recordings were performed using a Multiclamp 700B amplifier (Axon Instruments, Molecular Devices) and a Digidata 1550B digitizer (Axon Instruments, Molecular Devices) coupled with a computer equipped with pClamp11 software (Molecular Devices). Thin-walled borosilicate glass with filament (O.D.: 1.5mm, I.D. 1.1mm, 10cm length, fire polished, Sutter Instrument) was used for preparing recording pipettes using a horizontal puller (P-97, Sutter Instruments). The recording pipettes (4 to 5.0MΩ) were filled with internal solution containing the following: 120mM K-gluconate, 5mM NaCl, 2mM MgCl_2_, 0.1mM CaCl_2_, 10mM Hepes, 1.1 EGTA, 4mM Na_2_ATP, 0.4 Na_2_GTP, and 15 Na_2_Phosphocreatine (adjusted pH to ∼7.3 with HCl and osmolarity to ∼290 with sucrose). After establishing the whole-cell recording configuration, pipette capacitance was compensated before recording. Series resistance (Rs) was monitored (but not compensated), and only recordings with Rs less than 20 MΩ were included in this study. Current clamp bridge balance was adjusted prior to each action potential (AP) family recording. Recordings were low-pass filtered at 3 kHz and sampled at 10 kHz. Membrane potential values were not corrected for liquid junction potential. All recordings were conducted at room temperature. To measure intrinsic neuronal properties, resting membrane potential (RMP) was measured using a gap-free current-clamp recording without current injection for 10 seconds. Other intrinsic membrane properties and firing patterns were assessed in current-clamp mode using step current injections at the RMP or with the membrane potential held at -70mV. For step current injections, 500-ms pulses were applied beginning at -25 pA and increased in 5-pA increments up to a maximum depolarizing current of 225 pA. Input resistance was determined from the steady-state voltage responses to a series of 500-ms hyperpolarizing current steps (-25 to 0 pA, 5-pA increments) and calculated as the slope of a linear least-squares fit to the resulting current-voltage (I-V) relationship. The rheobase was defined as the minimal current step required to evoke at least one action potential (AP). The voltage threshold (mV) for AP initiation was defined as the membrane potential at which dV/dt exceeded 10% of its maximal value, relative to a baseline dV/dt measured 2 ms before the AP peak. AP amplitude was calculated as the difference between the voltage threshold and peak membrane potential. AP half-width was measured at half-maximal amplitude from the first evoked AP. AP onset latency was defined as the time (ms) between the start of the current injection and the threshold of the first AP. The fast afterhyperpolarization (fAHP) following the first evoked AP was calculated as the difference between the voltage threshold and the minimum membrane potential reached during the fAHP. Voltage sag (Vsag) was measured during a 500-ms hyperpolarizing current step (-25 pA) Percentage Vsag was calculated as: 100 x (ΔVpeak – ΔVss)/ ΔVpeak, where ΔVpeak is the peak voltage deflection and ΔVss is the steady-state voltage deflection.

For defining limitations of optogenetic parameters, we dissociated and cultured myenteric plexus preparations from *Pdyn*^ChR2-EYFP^ mice. Light-evoked action potentials were recorded in current-clamp mode in Pdyn-lineage neurons expressing ChR2-EYFP. A 470nm LED delivered 20 ms light pulses (20mW/mm2) through the objective (40x) to assess neuronal responsiveness. To establish parameters for in vivo use, neurons were stimulated for 20s at 1-15Hz. Spike probability was calculated as the ratio of evoked action potentials to total light pulses. Trials were repeated after a 2-minute recovery to confirm reproducibility and monitor opsin desensitization.

### Optogenetic activation of *ex vivo* colons

Mice were anesthetized with isoflurane and then euthanized by decapitation. The abdomen was dissected, and the colon was transected distally to the cecum and proximally to the rectum. The colon was pinned with minutien pins on a Sylgard Dish with the distal end open to allow expulsion of natural stool pellets. Colons were bathed in bubbled Krebs solution. A Logitech C615 1080pHD web camera accessed via Windows camera app was used to take video at 30FPS. A fiber optic (200µm diameter core; BFH48-200-Multimode, NA 0.48, Thorlabs) was connected to an 470µm LED and fixed about 3mm above the colon. Photostimulation had a maximum intensity of 18mW/mm^2^. Stimulation of 10Hz with 10ms pulse width for 20 seconds was delivered using a stimulator every 3 minutes for 25minutes. Videos were converted to spatial temporal maps using a custom MATLAB script (Mathworks). A propagating contraction was defined as a change in width of the colon that resulted in positional change of the natural stool pellet. The number of propagating contractions was quantified. The time from the onset of the light pulse and the initiation of the movement of a stool pellet was calculated as the time difference.

### Implantation of biocompatible LED

Mice were anesthetized with 2-4% isoflurane and eye lubricant was applied. Mice were kept warm with heating pad. Abdominal hair was shaved. Skin was numbed with bupivacaine and then sterilized with betadine. Then, a small vertical incision was made through the skin. The cecum was identified through the translucent fascia. About 1-2mm incision was made into the fascia. Then, a biocompatible 470 µm light emitting diode, LED (Neurolux unilateral brain device), was manipulated to allow passage of light into the peritoneum over the proximal colon and distal to cecum. The coil of the device was glued with vet approved cyano-acrylate on the fascia directly under the skin. The fascia was closed with nylon sutures. The skin was closed with staples, and Buprenorphine SR was administered subcutaneously. Staples were removed on day 7 post operatively. Behavioral and stool assessments occurred 7 days after staple removal.

### *In vivo* optogenetics

Mice were acclimated for 30 minutes to the chamber lined with wires attached to antenna, which was programmed using Neurolux program. The stimulator was programmed to give 10Hz pulses with 10ms pulse width for 1 minutes every 5 minutes (thus 4 minutes of recovery between stimulations). The stool produced during acclimation was cleared. Mice were filmed with a Sony HDR-CX190 handycam for 30 minutes. Total stools were counted at the end of 30 minutes. Total duration of not moving was timed after the 5^th^ stimulation. Orbital tightening was quantified when the mouse was facing the camera using the scale defined in the literature^42^ by an experimenter blinded to the genotype and treatment condition of the mice.

### Statistics

All statistics were performed in GraphPad Prism 11. Normally distributed data was compared using unpaired two tailed t tests and nonparametric data was compared using Welch’s t two tailed test.

## Acknowledgements

We appreciate experimental input and support from Drs. Rodney Newberry, Ellen Schill, and Vijay Samineni. We appreciate suggested edits from Dr. Phillip Tarr.

## Author contributions

D.J.V conceptualized the project, conducted experiments, and wrote the manuscript, J.L., P.G., B.J., J.W., and A.D. conducted experiments and provided edits to the manuscript, R.W.G. helped to conceptualized experiments and edited the manuscript.

## Conflicts of Interest

Dr. Robert W Gereau IV is a co-founder of Neurolux. Funding: This work was supported by NIH T32 DK077653 to D.J.V. and NIDDK R01DK116178 to R.W.G.

## Data Availability

The data presented in this study are available upon request from the corresponding authors.

## Notes

### Competing Interest Statement

The authors have declared no competing interest.

