## Supplemental Figures for "The Critical Role of *Pdyn*-Lineage Enteric Neurons in Colonic Motility and Visceral Interoception"

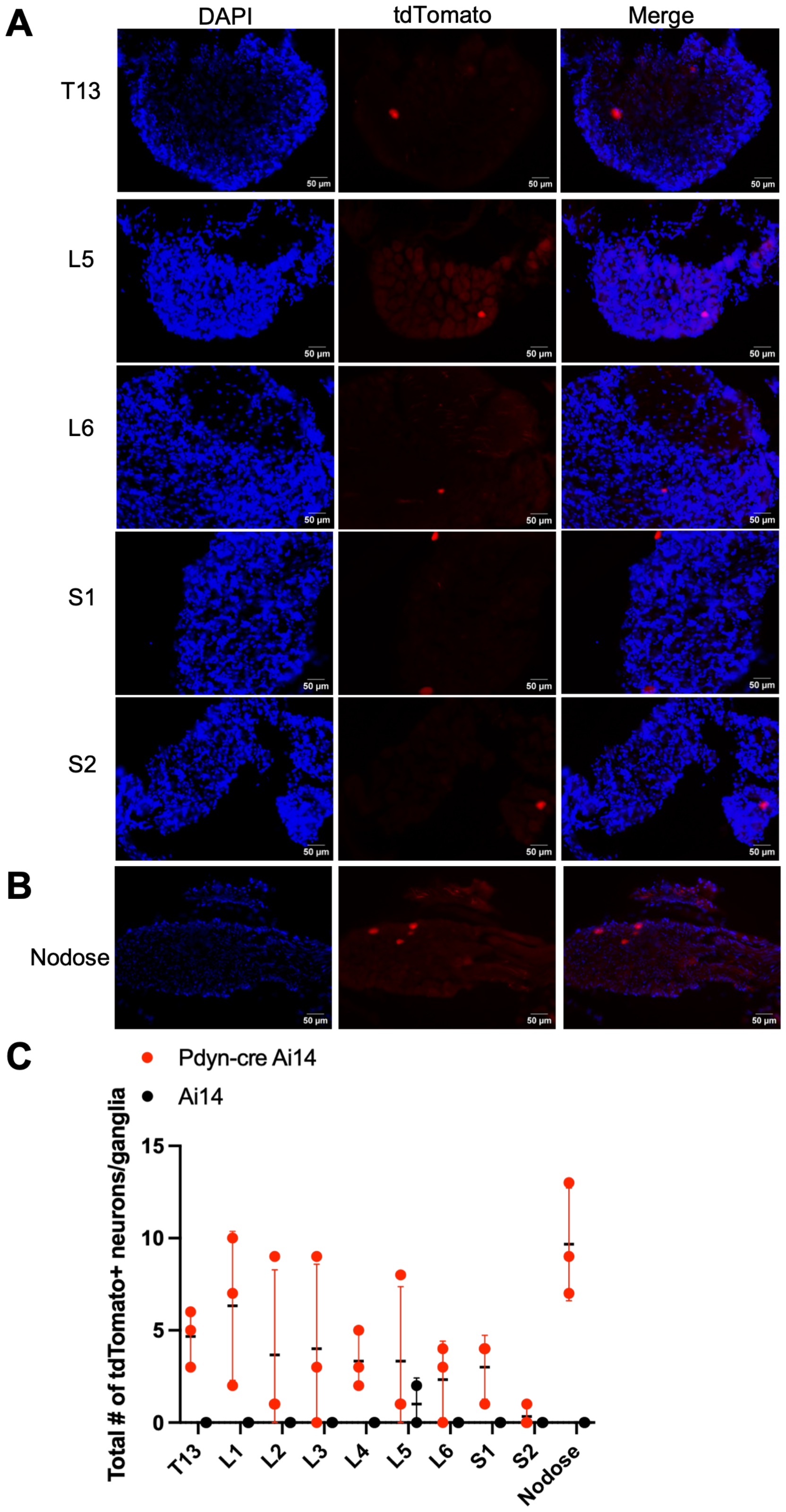

A
DAPI
tdTomato
Merge
T13
L5
L6
S1
S2
50 µm
B
Nodose
C
Pdyn-cre Ai14
Ai14
Total # of tdTomato+ neurons/ganglia
15
10
5
0
T13
L1
L2
L3
L4
L5
L6
S1
S2
Nodose

**Figure S1. *Pdyn*-expressing neurons of dorsal root ganglia and nodose ganglia of mice**

**A**. Representative images of cryosections of dorsal root ganglia at spinal levels T13-S2 from $Pdyn^{tdTomato}$ mice stained with DAPI. **B**. Representative images of cryosections of nodose ganglia from $Pdyn^{tdTomato}$ mice stained with DAPI. **C**. The total count of tdTomato expressing cells in each ganglia were quantified from $Pdyn^{tdTomato}$ (n=3) and control mice (n=2), which harbor the Ai14 allele but no Cre recombinase. Bars represent median and standard deviation.

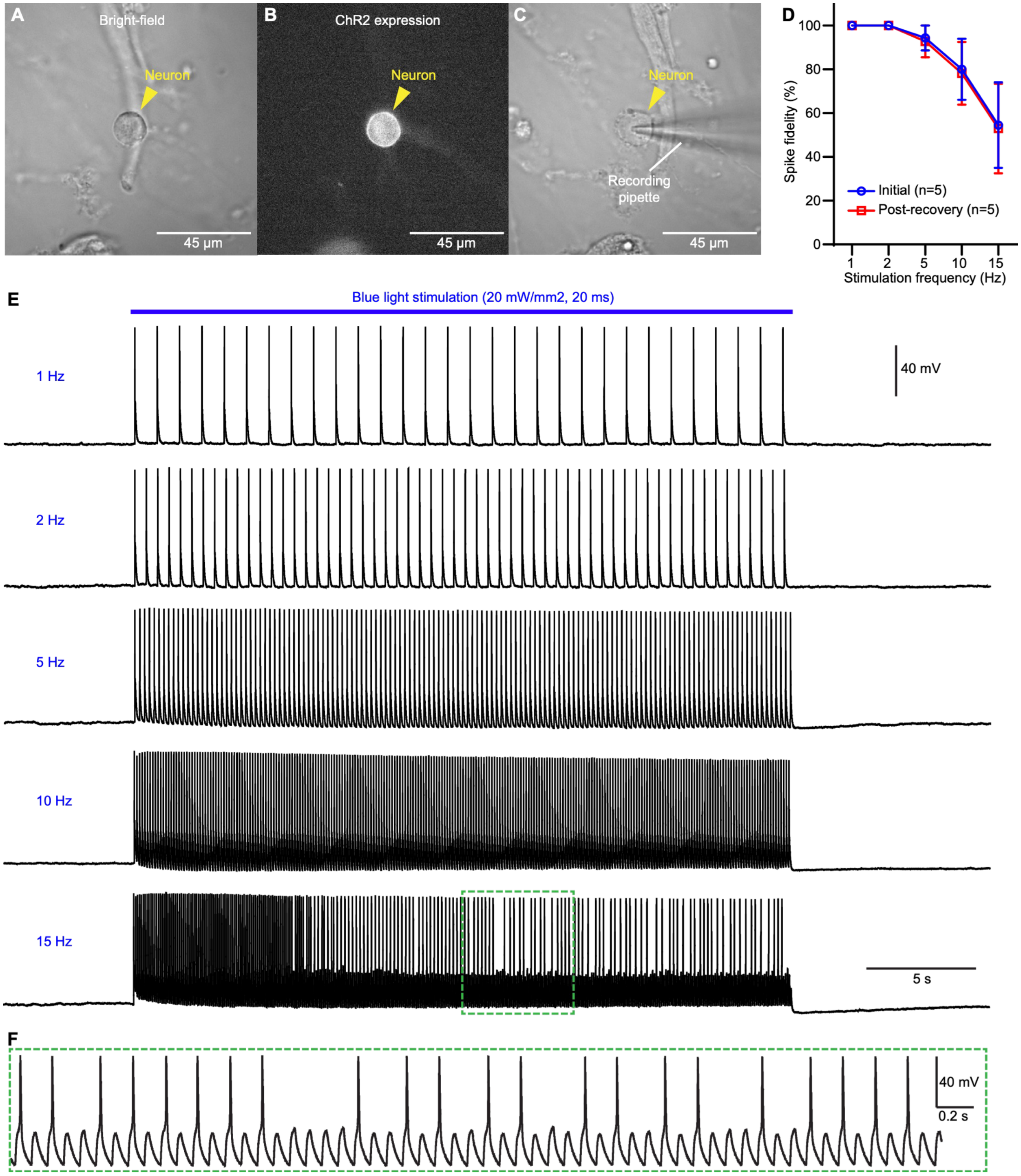


**Figure S2. Optogenetic activation of cultured *Pdyn*-expressing myenteric cells.**

**A-C**. Representative images of a cultured myenteric cells from *Pdyn*$^{ChR2\text{-}EYFP}$ mice, which are offspring of *Pdyn*-cre mice bred to Ai32 (ROSA26$^{LSL\text{-}ChR2\text{-}EYFP}$) mice, during whole-cell patch-clamp recording. The target cell is shown under bright-field (**A**), EYFP fluorescence indicating ChR2 expression (**B**), and with the recording pipette in place (**C**). Scale bars: 45 μm. **D**. Spike probability across different stimulation frequencies (1-15Hz) using 470nm blue light (20ms pulse width, 30 s duration). Data represent mean ± SEM for initial recordings (blue, n = 5) and post-recovery trials (red, n = 5). **E**. Representative whole-cell current-clamp traces showing action potentials evoked by blue light stimulation (blue bar, 20mW/mm2) at frequencies of 1, 2, 5, 10, and 15Hz in cultured myenteric cells with *Pdyn* expression in their lineage. **F**. Representative whole-cell patch-clamp trace during 15Hz light pulse showing unreliable action potential generation.

| Parameter | Value (mean ± SEM) |
| --- | --- |
| **Ithreshold (pA)** | 27.0 ± 5.0 |
| **Vthreshold (mV)** | -13.9 ± 1.9 |
| **Onset (ms)** | 58.9 ± 8.6 |
| **fAHP (mV)** | -20.0 ± 0.9 |
| **APamplitude (mV)** | 45.7 ± 3.7 |
| **APD50 (ms)** | 4.6 ± 0.4 |
| **Vsag (%)** | 1.6 ± 0.5 |
| **sAHP (mV)** | -3.4 ± 0.5 |
| **Cm (pF)** | 17.3 ± 3.9 |
| **n** | 10 |

**Table S1**. Electrophysiological properties of myenteric neurons that express *Pdyn* in their lineage.
